# A Strategy to Identify Endogenous Ligands for a Protein

**DOI:** 10.64898/2026.11.20.768119

**Authors:** J. Conrad Epps, D. Ryan Bhowmik, Safal Shrestha, Christopher Esselman, Samiksha Katiyar, Emory Gandy, Natarajan Kannan, Arthur S. Edison

## Abstract

Protein-metabolite interactions are essential to numerous metabolic processes and underly many disease mechanisms. However, detecting these interactions is a challenge due to the transient nature of protein-metabolite associations. Here, we employ an untargeted functional metabolomics screening strategy to uncover low-affinity interactions between an entire metabolome and a kinase. We first utilize high-performance liquid chromatography (HPLC)-guided fractionation to separate a whole-metabolome extract into multiple fixed-time fractions. A metabolite fraction library (mFL) of one-dimensional (1D) proton (^1^H) NMR spectra is then generated from these fractions. After adding the protein to the same fractions, saturation transfer difference (STD) NMR is then performed on each to detect weak protein-metabolite interactions and develop an STD library. Using the binding resonances from the STD library as a guide, the identity of the binding ligand can be determined. We demonstrate this strategy using frutosamine-3-kinase (FN3K) to identify and validate both known and novel binders. This workflow serves as a practical and simple way to screen an organism’s metabolome for endogenous effector molecules, further strengthening our knowledge of protein-metabolite interactions and the functional metabolome.

**Abstract Graphic:** 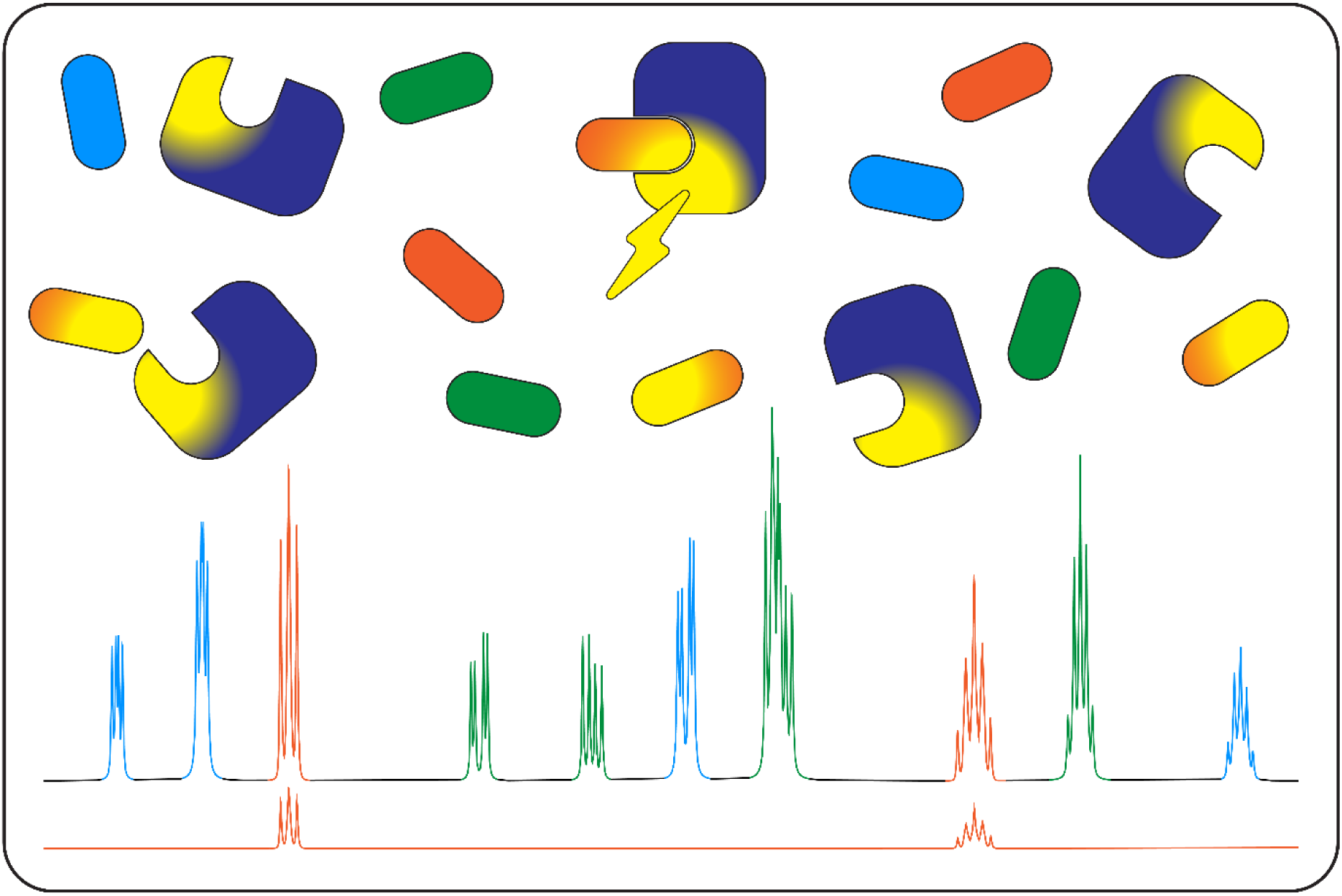

## Introduction

Metabolites bind to proteins and other macromolecules as substrates, inhibitors, and allosteric effectors. The fields of proteomics^1^ and metabolomics^2^ have matured to a point where extensive details within each biological omics level can be obtained. However, much of biology depends on the interactions between different biological levels, such as proteins and metabolites, and methodologies to investigate these interactions on a systems biology scale are lacking.^3^

Much work has been dedicated to defining the metabolome in a global sense through untargeted metabolomics.^2, 4^ Connecting metabolites to specific activities and functions is challenging, and methods to do this are often specific to a phenotype or disease of interest.^5–8^ For protein-metabolite interactions in particular, identifying endogenous ligands for a specific target protein remains difficult.

The pharmaceutical industry has long-used saturation transfer difference (STD) nuclear magnetic resonance (NMR) spectroscopy to screen large libraries of ligands for their ability to bind to a protein target of interest.^9, 10^ STD NMR is a ligand-observed approach, which uses sub-stoichiometric amounts of a protein and relies on rapid exchange rates between the free and bound states of the ligand.^11, 12^ As such, STD NMR is a useful technique to assess binders with weak-to-moderate binding affinities.^13, 14^

Nikolaev et al. previously used STD NMR experiments to identify known interactions between a set of proteins and metabolites.^15^ This points to the feasibility of using STD NMR to screen for weak binding interactions between metabolites and enzymes. Diether et al. utilized a similar ligand-observed NMR method to identify 98 previously unknown binding interactions between 55 known metabolites and 35 central carbon metabolic enzymes in *Escherichia coli*.^16^ This study demonstrates the large gap in knowledge in this area, even for one of the most well-studied organisms in science. While this study utilized a known set of proteins and metabolites, to our knowledge, STD NMR and other ligand-binding NMR experiments have not been reported for use in the untargeted screening of an entire metabolome with a specific protein.

Our study expands the application of STD NMR to evaluate binding interactions between a small-molecule protein kinase and a whole metabolome. We used fructosamine-3-kinase (FN3K), which is known to interact with nicotinamide-containing cofactors of central metabolism, such as NAD^+^.^17, 18^ We identified a previously known interaction involving NAD^+^ and 35 novel interactions. We then tested a subset of 6 STD NMR binders and validated their interactions using thermal stability assays.

## Results

### Synthetic nicotinamides bind to FN3K

We used STD NMR to measure four synthetic nicotinamide cofactors mixed with FN3K. The resulting STD difference spectra shown in **Figure 1** are consistent with the previous results regarding apparent strength of interaction: The NAD^+^/NADH redox pair showed the most intense interactions with the protein as compared to NADP^+^/NADPH. Full binding data are shown in **SI Figure 1**.

**Figure 1.**
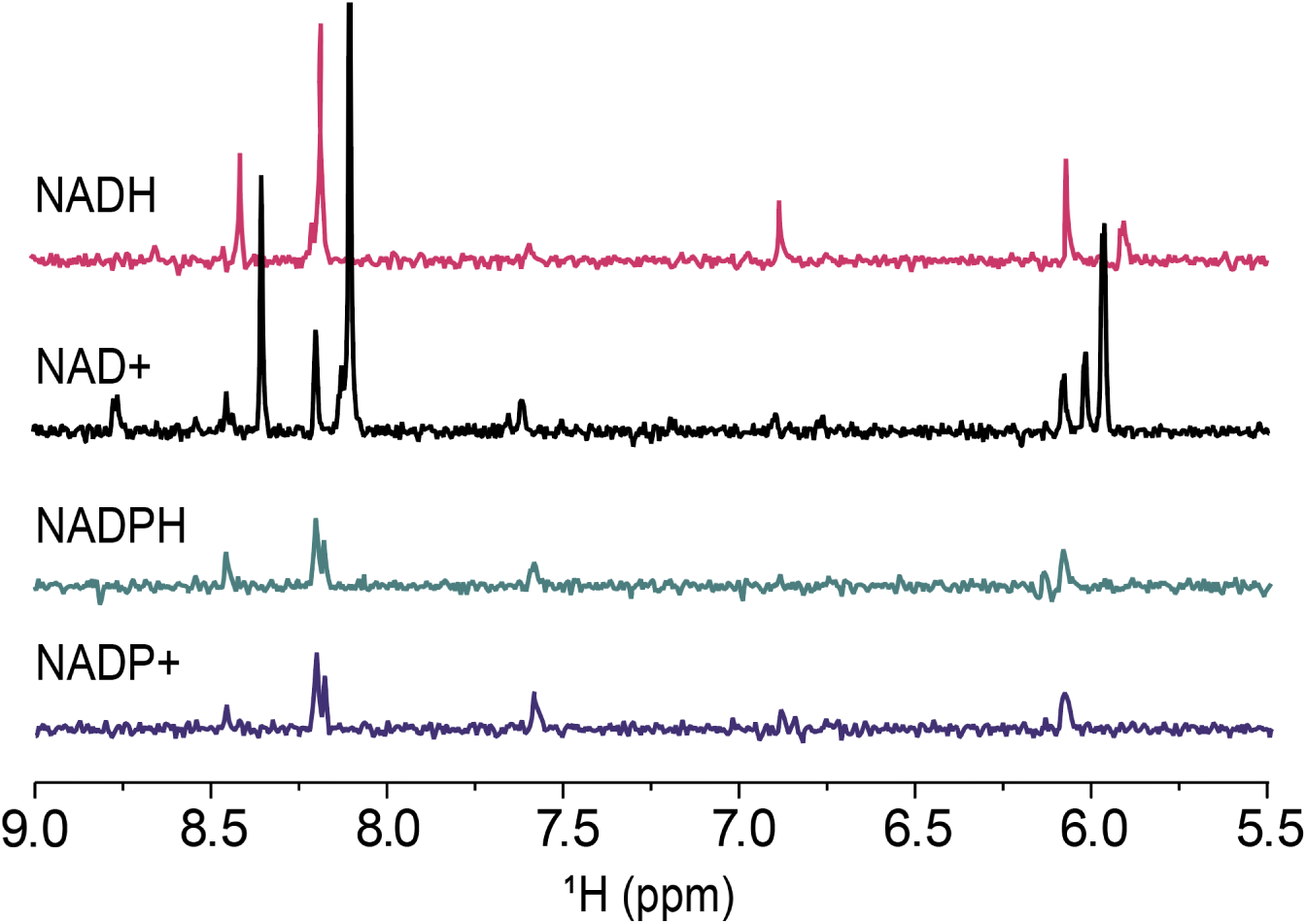
| FN3K binding to nicotinamide cofactors. STD NMR difference spectra of individual NAD compounds shown.

### STD NMR selects binding compounds from a mixture of metabolites

After validating that synthetic nicotinamide compounds bind to FN3K, we investigated an experimental metabolite mixture. We used a polar metabolome extract from *Caenorhabditis elegans* and acquired an STD NMR spectrum of the mixture with FN3K added. **Figure 2** shows a set of downfield resonances similar to those seen in the synthetic nicotinamide experiments. Additionally, several unexpected resonances appeared, presenting the possibility of novel ligand binders for the protein.

**Figure 2.**
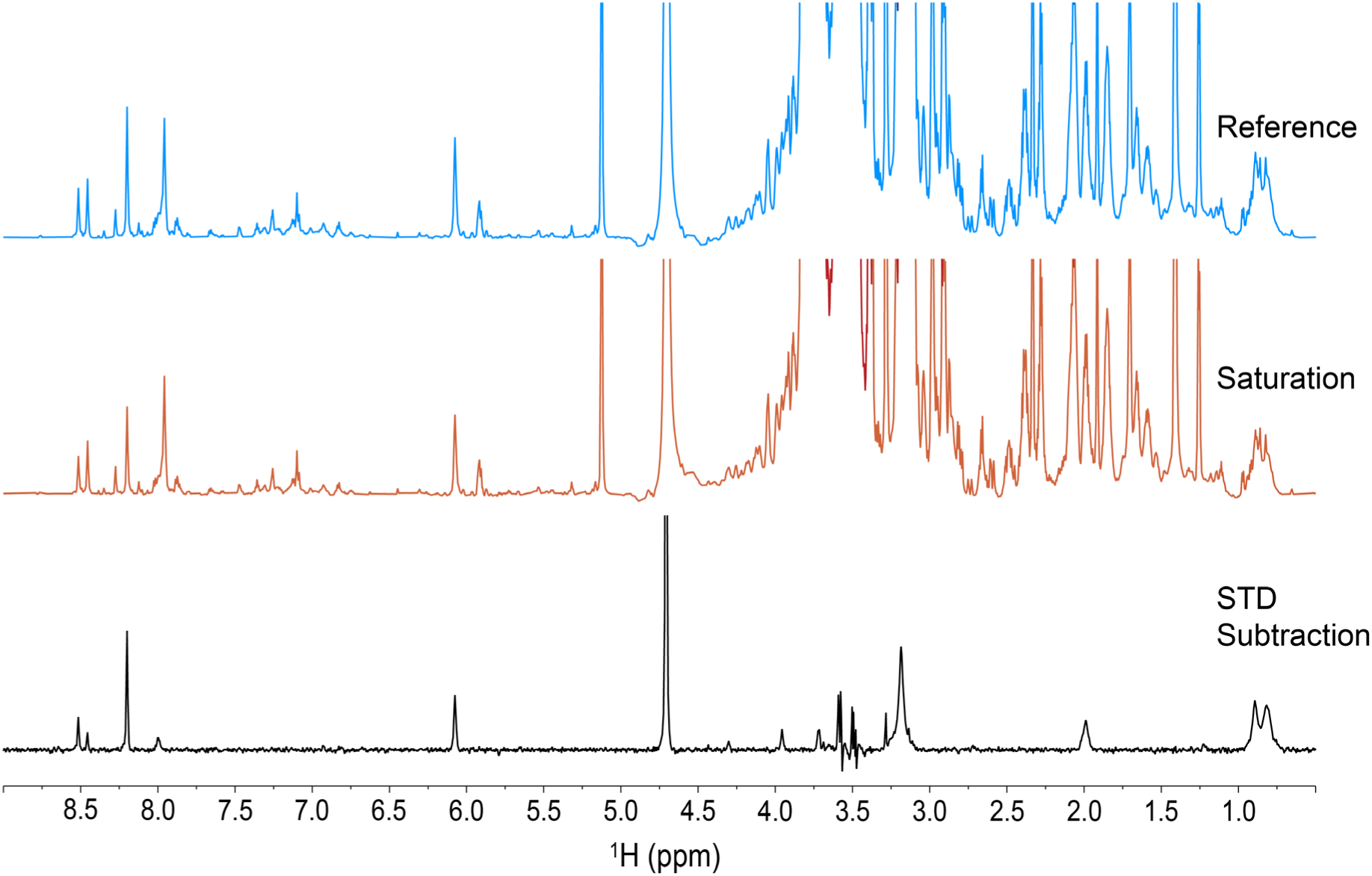
| FN3K screens binding metabolites from an entire metabolome by STD NMR. Complete STD NMR dataset: reference spectrum of mixture in blue, saturated spectrum in orange, and difference spectrum in black.

### Fraction libraries enable non-targeted annotation of a complex metabolite mixture

To quantify and identify the compounds present in the *C. elegans* metabolome mixture, we fractionated the sample using semi-preparative hydrophilic interaction liquid chromatography (HILIC). On each of the 140 fractions, we collected one-dimensional (1D) proton (^1^H) NMR spectra to generate a metabolite fraction library (mFL) as shown in **Figure 3a**. The fractionation reduced peak overlap and enabled the correlation of resonances across eluting fractions. Using FLAnalyzer in a non-targeted manner, as originally published, 105 metabolite basis set (mBS) elements were generated, representing likely metabolites in the library based on high correlation values (>0.98).^19^ Of the 105 mBS elements, 30 matched the COLMAR^20^ database with high confidence, 14 matched with lower confidence, and 61 had no match. (**SI Table 1**).

**Figure 3.**
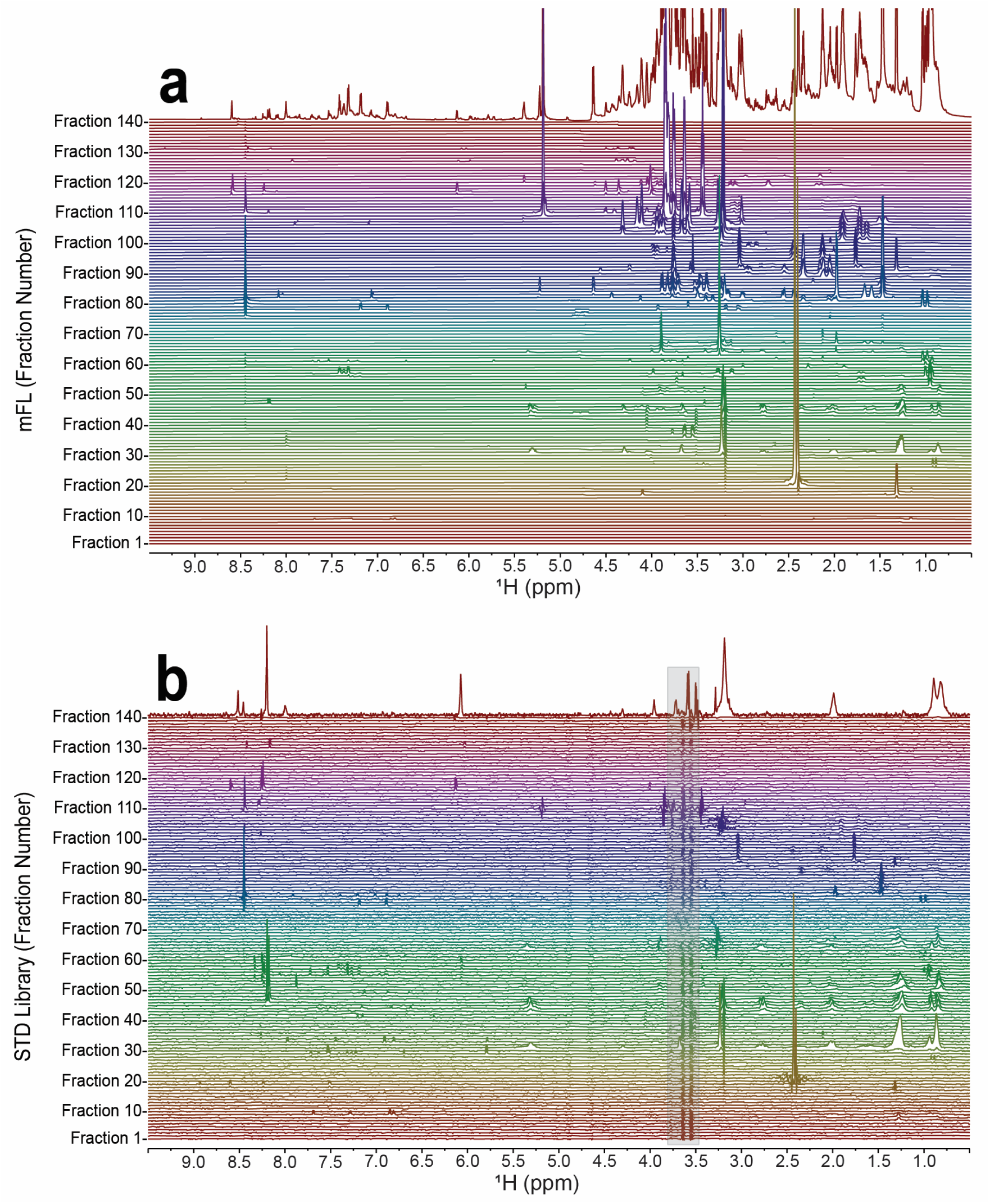
| [**a**] 1D ^1^H spectra of mFL fractions derived from an extract of *C. elegans.* The top trace is the unfractionated mixture. [**b**] The same metabolite library fraction samples screened with *A. thaliana* FN3K by STD NMR (STD library) and shown as difference spectra. The top trace represents the STD subtraction of the unfractionated mixture binding experiment. Glycerol present from protein addition is boxed in grey.

### STD NMR enables targeted annotation of binding interactions within a whole metabolome

By adding FN3K to each mFL sample, an STD library was generated. The STD experiments, combined with chromatographic separation, revealed a limited number of distinct peaks, as shown in **Figure 3b**. Each peak shown represents a potential binding interaction. Regional expansions for **Figure 3** are shown in **SI Figure 2**. To identify the binding compounds, STD library peaks were used as a guide when selecting driver peaks in the mFL using FLAnalyzer. **Figure 4** displays an overlay of the STD NMR difference spectra with their respective basis element ^1^H spectra for all STD peaks shown. A similar up-field expansion is shown in **SI Figure 3**, and the full STD library, including all identified compounds, is displayed as overlays in **SI Figure 4**. Targeted annotation of the STD library for basis element generation identifies both known and unknown compounds from the complex metabolome. The targeted mBS consists of 36 basis set elements. After database matching, 21 basis elements match COLMAR with high confidence, 1 element has a lower confidence match, and 14 have no obvious matches (**SI Table 2**). The method for generating the targeted basis set is outlined in **SI Figure 5**. Cosine similarity was performed to determine how well the non-targeted mBS matched the targeted mBS. High similarity scores of >0.90 were seen for 21 out of the 36 basis set elements, as shown in **SI Table 3**.

**Figure 4.**
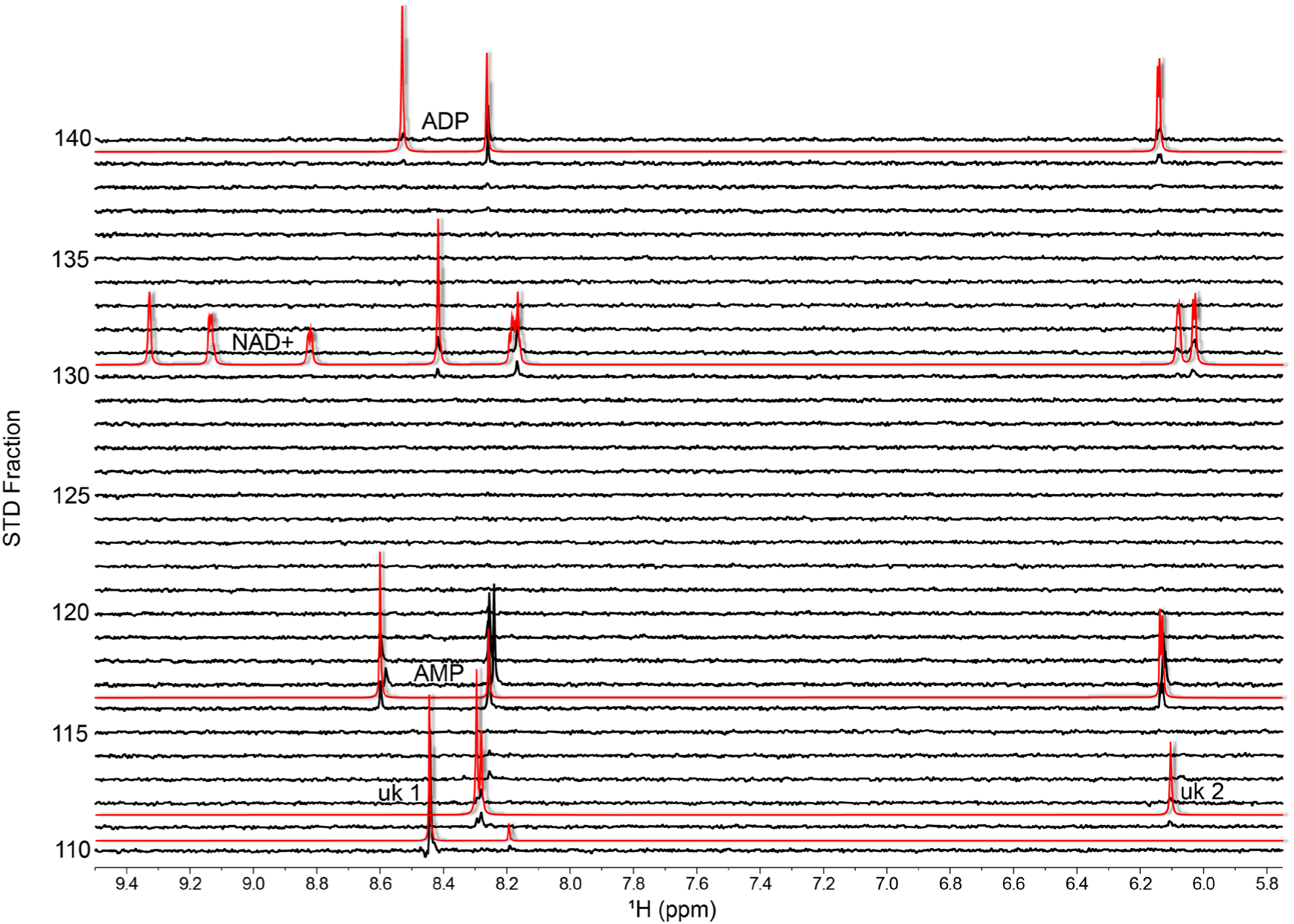
| STD NMR screen of the *C. elegans* mFL. STD NMR difference spectra (black) for each fraction in a subset of the library are overlaid with 1D ^1^H spectra of basis set elements (red), isolated using STD peaks as a guide. Unknown basis set elements marked as “uk”.

**Figure 5.**
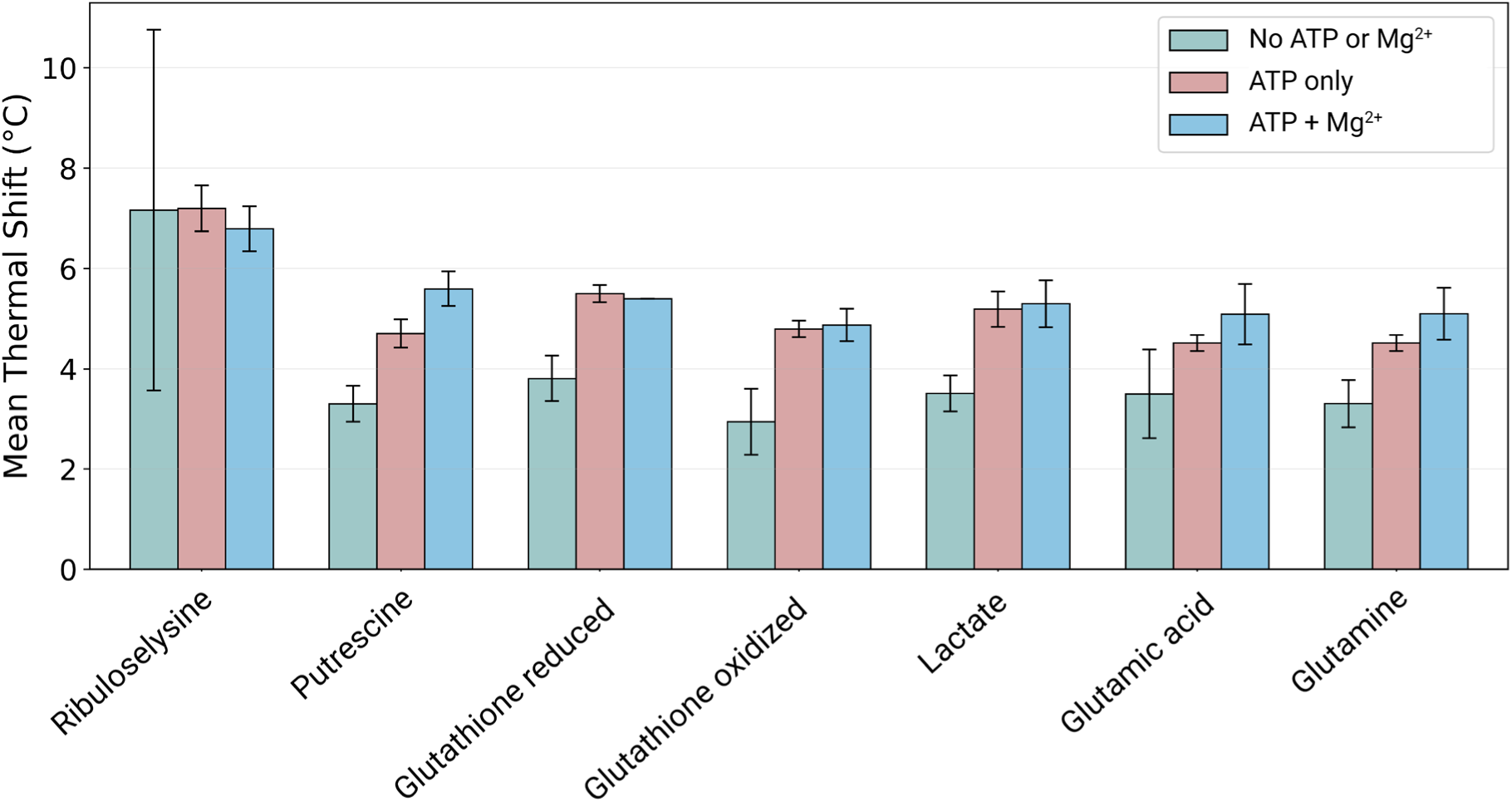
| Mean thermal shift (ΔT) for each compound with FN3K. Each compound was treated to three different conditions: No additional cofactors (green), ATP only (salmon), and ATP + Mg^2+^ (blue).

### Validating a selection of putative compounds as FN3K binders

From the 21 matched basis elements, six potential interactors were selected for binding validation by thermal shift assay. The selected compounds, putrescine, glutathione (oxidized and reduced forms), lactate, glutamic acid, and glutamine, were previously implicated in pathway enrichments for joint transcriptomic and metabolomic studies of FN3K knockout cell lines. Glutathione metabolism was upregulated. Putrescine, glutamine, and lactate were also specifically identified in association mapping of differentially expressed genes and differentially accumulated metabolites in the context of vitamin and cofactor metabolisms.^18^ Ribuloselysine serves as a positive control, as ribulosamines are known to be FN3K binders.^21^ Each selected compound is seen to stabilize FN3K (**Figure 5**). Additionally, the strength of binding shows an increasing trend as ATP and Mg^2+^ are added to the binding mixture. The strength of the nicotinamide binding has been previously shown to increase in a metal-dependent manner.^18^

## Discussion

Using metabolite fraction libraries and STD library screening we have outlined a simple method to screen a whole metabolome for protein-metabolite interactions. This approach uses one HPLC fractionation, and two NMR runs of the same sample set. The nondestructive nature of NMR spectroscopy as an analytical technique is essential for this approach. This methodology can be applied to any number of proteins or other macromolecules, provided the binding affinities are weak to moderate.

With STD, it is important to try to distinguish nonspecific from specific binding. STD NMR can generate false positives due to signal accumulation from multiple binding sites across the protein, substrate amalgamation, and the relative strength of the binding interactions themselves.^22^ As shown for NAD^+^, conducting binding experiments with orthogonal methods, such as thermal stability assays with synthetic compounds, is important to rule out false positives.

In contrast to the four nicotinamides previously implicated in FN3K metabolism, only NAD^+^ was detected from the *C. elegans* extract. We might expect to see the other nicotinamides based on the binding patterns from **Figure 1** and the strong intensities seen for the synthetic NADH/NAD^+^ redox pair. *In vivo*, the intercellular concentration of NADH is low compared to NAD^+^.^23^

After chromatographic separation, more peaks appear in the STD library than in the initial mixture STD experiment, revealing a larger set of binding interactions. We suspect that some part of these interactions could be due to the removal of competitive binding by separating the mixture.

Here, we have demonstrated a simple way to screen a protein for endogenous metabolite binding, which could help elucidate new mechanisms of enzyme regulation.

## Methods

### Data Availability

Raw NMR data and basis set elements can be found on the NAN resource connector.^24^ FN3K synthetic nicotinamide experiments: usnan.org/ark:/83454/c13d5ec807-a4e1-45f6-9165-d22d6f20c25a.1 FN3K STD in unfractionated *C. elegans* mixture: usnan.org/ark:/83454/c143c10d4c-0090-40e6-89ec-66247d2f35f6.1 mFL 1D data and supplemental non-targeted basis set elements: usnan.org/ark:/83454/c1141c7e1e-c9f8-42c9-9ced-90f0e44a2850.1 STD library data and supplemental targeted basis set elements: usnan.org/ark:/83454/c19723b772-2c39-42cd-b247-b767da00d99d.1

### *C. elegans* Growth and Maintenance

The *C. elegans* reference strain CGC1 (PD1074) available at the Caenorhabditis Genetics Center (University of Minnesota) was used to generate worm stocks. Worms were grown and harvested after a previously developed protocol for the culture of large-scale mixed-stage *C. elegans* populations.^25^ After culturing, nematodes were sorted by large particle flow cytometry into aliquots of 200,000 worms each, which were flash-frozen in ddH_2_O and stored at −80°C as part of the same protocol.^25^

### *C. elegans* Metabolite Extraction

100 cryovials (20 million nematodes) of the *C. elegans* stocks were lyophilized for 24 hrs at room temperature. Five 1.0 mm zirconia beads were added to each tube before bead-beating at 1800 rpm for 90 s with dry ice using a FastPrep-96 homogenizer. 300 µL LC/MS grade 80:20 MeOH/H2O was pipetted into each tube and vortexed for 20 seconds. The tubes were then centrifuged at 14,000 rpm at 4 °C for 30 min. The supernatant was transferred into a single tube and lyophilized at room temperature until dry. After storage at -80°C, the vacuum concentrate was reconstituted in 300 µL LC/MS grade 80:20 MeOH/H_2_O in the same tube it was dried in. The tube was spun down to sediment out all non-dissolved solids. The supernatant was then transferred to a 2 mL HPLC vial.

### HPLC Fractionation

The metabolite fraction library was generated by three 100 µL net extract injections at 25°C on an Agilent 1260 Infinity HPLC system using XBridge BEH Amide OBD Prep Column, 130 Å, 5 µm, 10 x 250 mm HILIC column. A consistently linear, 35 min gradient of 0.1% formic acid in H_2_O (A) and 0.1% formic acid in ACN (B) was used. 0-20 min was a 5% to 30% gradient of A, 20-30 min was a 30% to 50% gradient of A, and the run concluded with a with a 30-35 min gradient of 50% to 65% A. The entire gradient was run at a flow rate of 3.5 mL/min. Spectral data were collected using an Agilent Infinity Lab Single Quadrupole MSD. 140 equal, fixed-time fractions were collected, and each separate injection was added to the same vials during dry-downs. Between the second and third injections, and after the third, the vials were dried using a Centrivap vacuum concentrator. Dried fractions were stored at -80 °C.

### Protein Expression and Purification

AtFN3K gene cloned into pET-15b expression vector was provided by E. Van Schaftingen (Université Catholique of Louvain, Brussels, Belgium). *E. coli* BL21 (DE3) pLysS cells (Roche) were co-transformed with AtFN3K and GroEL (Addgene) for protein expression and purification. Co-transformed cells were cultured in 1L LB media containing 10% glycerol, 100 µg/mL ampicillin and 25 µg/mL chloramphenicol. Cells were induced at OD 0.6 with 250 mM IPTG and further grown at 16 °C for 18 hrs. After 18 hrs, cells were harvested and processed for protein purification using His-Pur Cobalt Resin (ThermoFisher scientific). Briefly, *E. coli* cells were lysed by sonication in 50 mM sodium phosphate pH 7.4, 300 mM NaCl, 15% glycerol and 1 mM PMSF (lysis buffer). Clarified cell lysate was passed through Cobalt-Resin column, washed 10x with lysis buffer and bound proteins were eluted with 300 mM imidazole. Purity of the eluted fractions was checked on SDS-PAGE gels and fractions containing AtFN3K at 35kDa were pooled for further purification by Size Exclusion Chromatography (SEC) using a HiLoad 16/1600 Superdex 200-pg column (Cytiva) equilibrated in SEC buffer (25 mM HEPES, pH 7.5, 300 mM NaCl and 10% glycerol). A total of 5ml AtFN3k purified from Cobalt-resin column was loaded and elution was performed at a flow rate of 1 mL/min. Purity of eluted fractions was further confirmed by SDS-PAGE gel followed by snap-freezing the pure protein at -80 °C which was then used to collect the NMR data.

### NMR Data Acquisition

NMR data were collected on an Oxford 800 MHz magnet with a Bruker Avance Neo console, 1.7 mm TCI cryoprobe, and SampleJet automatic sample changer. All 1D ^1^H NMR experiments were acquired using the “noesypr1d” pulse sequence. Data were collected with 32,768 points with 4 dummy scans and 64 scans each at 25°C

All STD NMR experiments were acquired using the “stddiffgp19.3” pulse sequence using an off-resonance frequency of -40 ppm and an on-resonance frequency of 0.2 ppm. Data were collected with 32,768 points with 4 dummy scans and 32 scans at 25°C.

### NMR Sample Preparation

All synthetic nicotinamide binding experiments were performed with 1 mM ligand against 25 µM protein (40:1 ligand-to-protein ratio) with 10 mM MgCl_2_, 200 mM NaCl, and 20 mM phosphate buffer. The STD experiment with FN3K in the total metabolite mixture was performed with extracted metabolites from 200,000 *C. elegans* with 25 µM *A. thaliana* FN3K. All individual library fractions and metabolite extracts were resuspended in 55 μL D2O buffer in 1.7 mm Bruker racks of 96 tubes. For STD library binding experiments, 25 µM protein was added to each sample in the library using a Bruker SamplePro NMR tube loading robot.

### NMR Data Processing

NMRPipe batch processing was used.^26^ Autophasing with zero-order phase correction, 2 Hz exponential line broadening, automatic first-order baseline correction, and DSS referencing were performed. 1D ^1^H data were reference deconvoluted with a line width of 1 Hz, line broadening of 1.5 Hz, and trapezoidal apodization of 75%.^27^ SAND modeling was conducted for 1D ^1^H experiments from a range of 10.3 ppm to -2.0 ppm.^27^ For STD experiments, the on- and off-resonance spectra were subtracted in the time domain to give the difference spectra.

### NMR Data Analysis

The MATLAB application “FLAnalyzer” (https://github.com/edisonomics/FLAnalyzer) was first used to generate the non-targeted mBS of 105 metabolites from highest to lowest intensity.^19^ The targeted mBS of 36 metabolites was generated using the STD peaks as a guide for setting driver peaks to begin basis element generation. To identify which elements were shared between the non-targeted and targeted libraries, cosine similarity was performed to determine which non-targeted elements best matched to targeted elements. After peak picking each individual basis set element, the chemical shift values for each were used as input into COLMAR 1D Query.^20^ The highest mean scoring matches were compared to BMRB reference spectra for each basis set element.^28^ Matches were assessed and ranked visually for their similarity. Matches are scored as with a confidence level of either high (all peaks matching), medium (some matching), or not determined (no peaks matching).

### DSF/Thermal Shift Assay

Differential scanning fluorimetry (DSF) was used to assess the thermal stability of *A. thaliana* fructosamine-3-kinase (AtFN3K) in the presence of selected metabolites. Reactions were prepared in a final volume of 50 µL containing 0.5 mg/mL AtFN3K and 1:100 SYPRO Orange in buffer containing 20 mM HEPES, pH 7.4, 150 mM NaCl, and 5% glycerol. The metabolites tested were putrescine, oxidized glutathione, reduced glutathione, lactate, glutamic acid, and glutamine. Ribuloselysine was included as a positive control. Each metabolite and ribuloselysine were tested at a final concentration of 50 mM.

AtFN3K was tested with each compound under three conditions: metabolite alone, metabolite with 50 mM ATP, and metabolite with 50 mM ATP and 25 mM MgCl_2_. Control reactions containing each compound in buffer without AtFN3K were included to assess background signals from the plate. The buffer volume was adjusted as needed to maintain a final reaction volume of 50 µL.

Samples were heated from 20 °C to 95 °C at a rate of 0.3 °C/min, and fluorescence was monitored using a StepOnePlus Real Time PCR System.

Thermal shifts (ΔT_m_) were calculated by subtracting the melting temperature (T_m_) of AtFN3K in the absence of metabolites from the T_m_ measured in the presence of each metabolite under the corresponding ATP and MgCl_2_ conditions. The resulting ΔT_m_ values were averaged for each condition and plotted as a grouped bar graph to compare metabolite-induced changes in AtFN3K thermal stability. The DSF data was processed and plotted using Matplotlib.

## Acknowledgements

We thank Dr. Amanda Shaver and Pam Kirby for culturing the *C. elegans* used in this study. Kennedi Clemmons helped with the preparation of NMR samples. John Grimes assisted with STD studies and NMR automation. Alex Eletsky helped with NMR processing. This work was supported by NIH grant R35GM148240 to ASE and R35GM148240-02S2 to ASE and JCE. Funding for NK (R35GM139656) is acknowledged.

---

**SI Figure 1.**
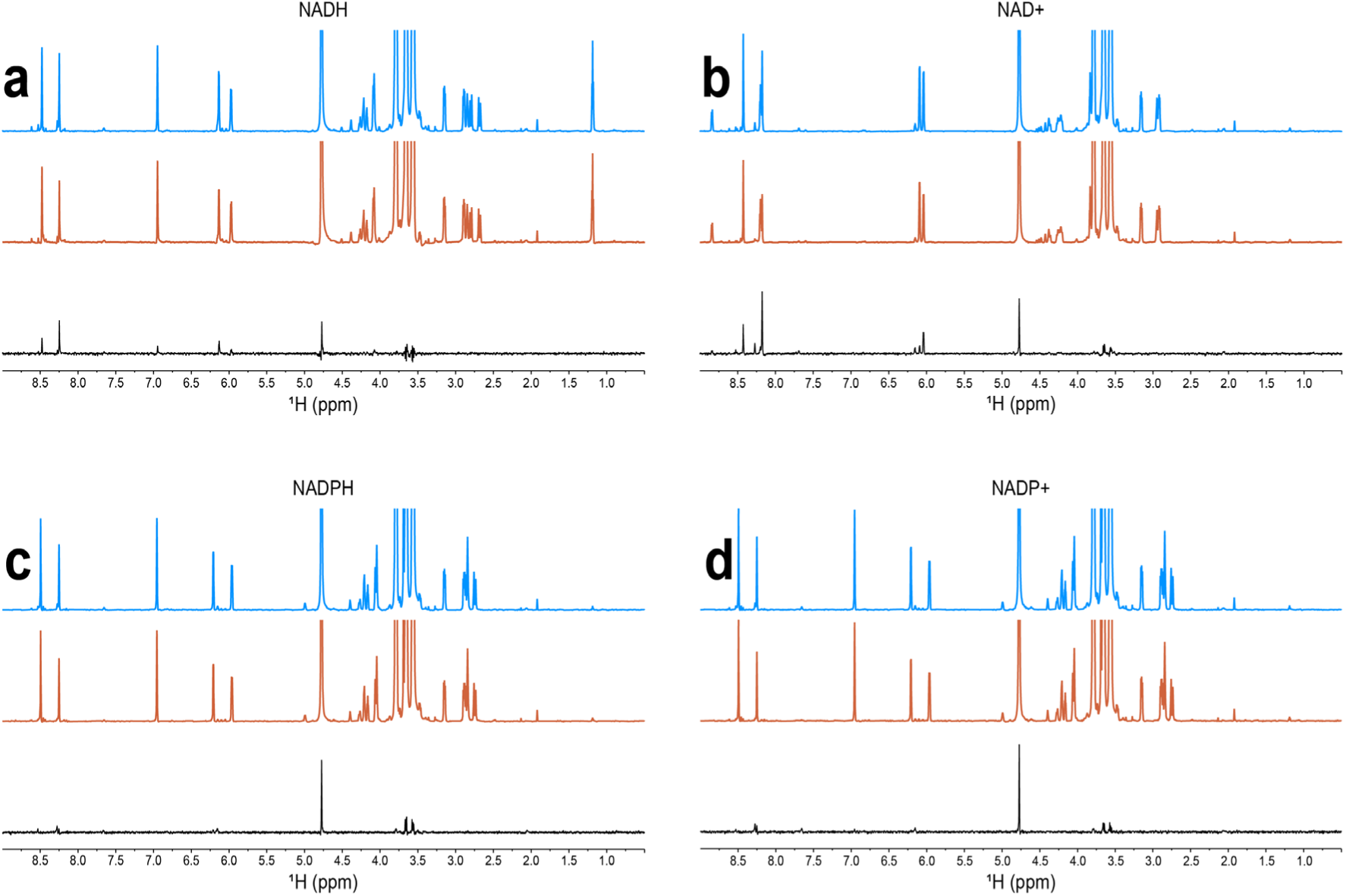
| *Arabidopsis* FN3K binding to nicotinamide cofactors. Complete STD NMR dataset: reference spectrum of mixture in blue, saturated spectrum in orange, and difference spectrum in black.

**SI Figure 2.**
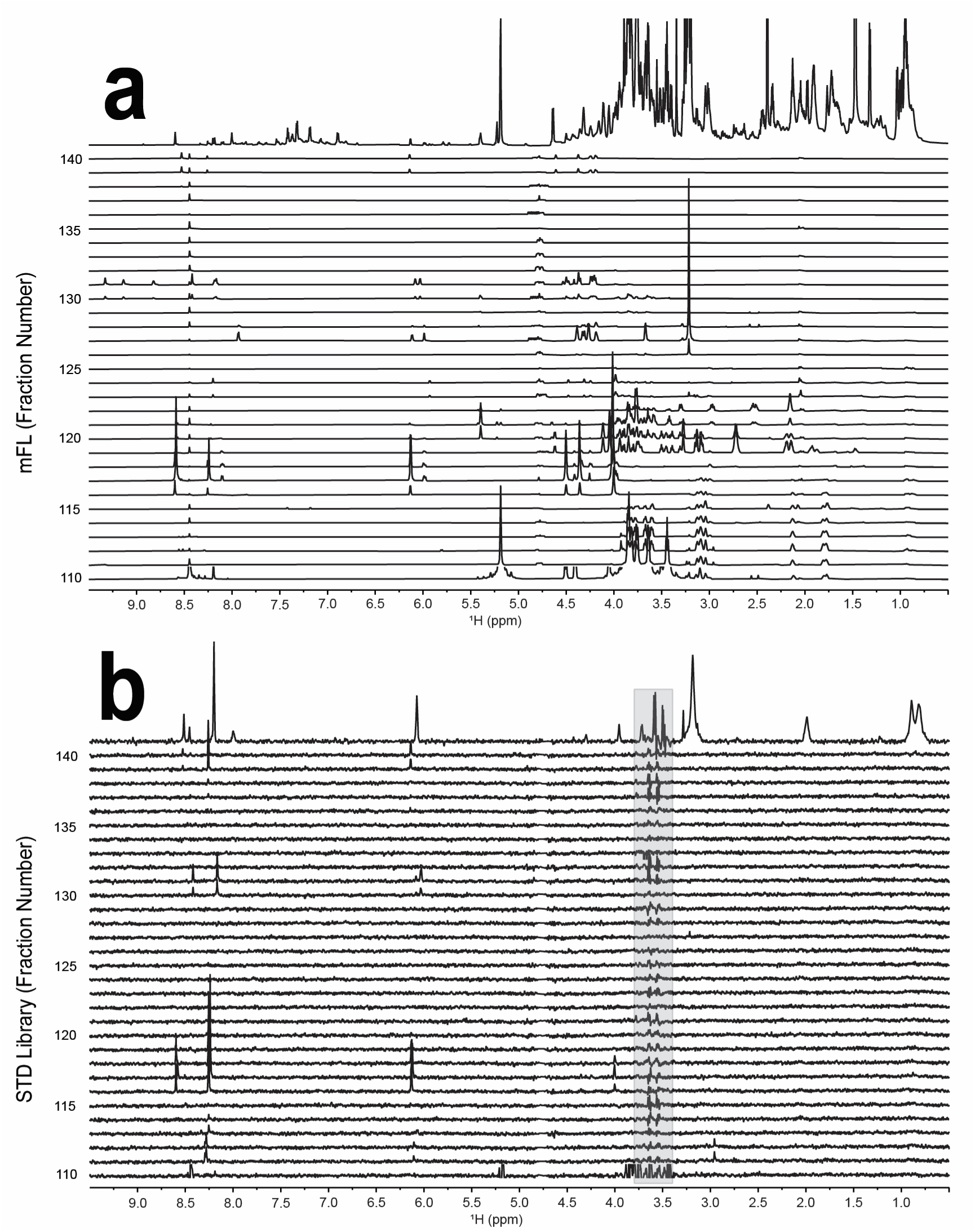
| [**a**] 1D ^1^H spectra of mFL fractions 116-131 derived from an extract of *C. elegans.* The top trace is the unfractionated mixture. [**b**] The same metabolite library fractions 116-131 screened with *A. thaliana* FN3K by STD NMR (STD library). All spectra shown are difference spectra. The top trace represents the STD subtraction of the unfractionated mixture binding experiment. Glycerol present from protein addition is boxed in grey.

**SI Figure 3.**
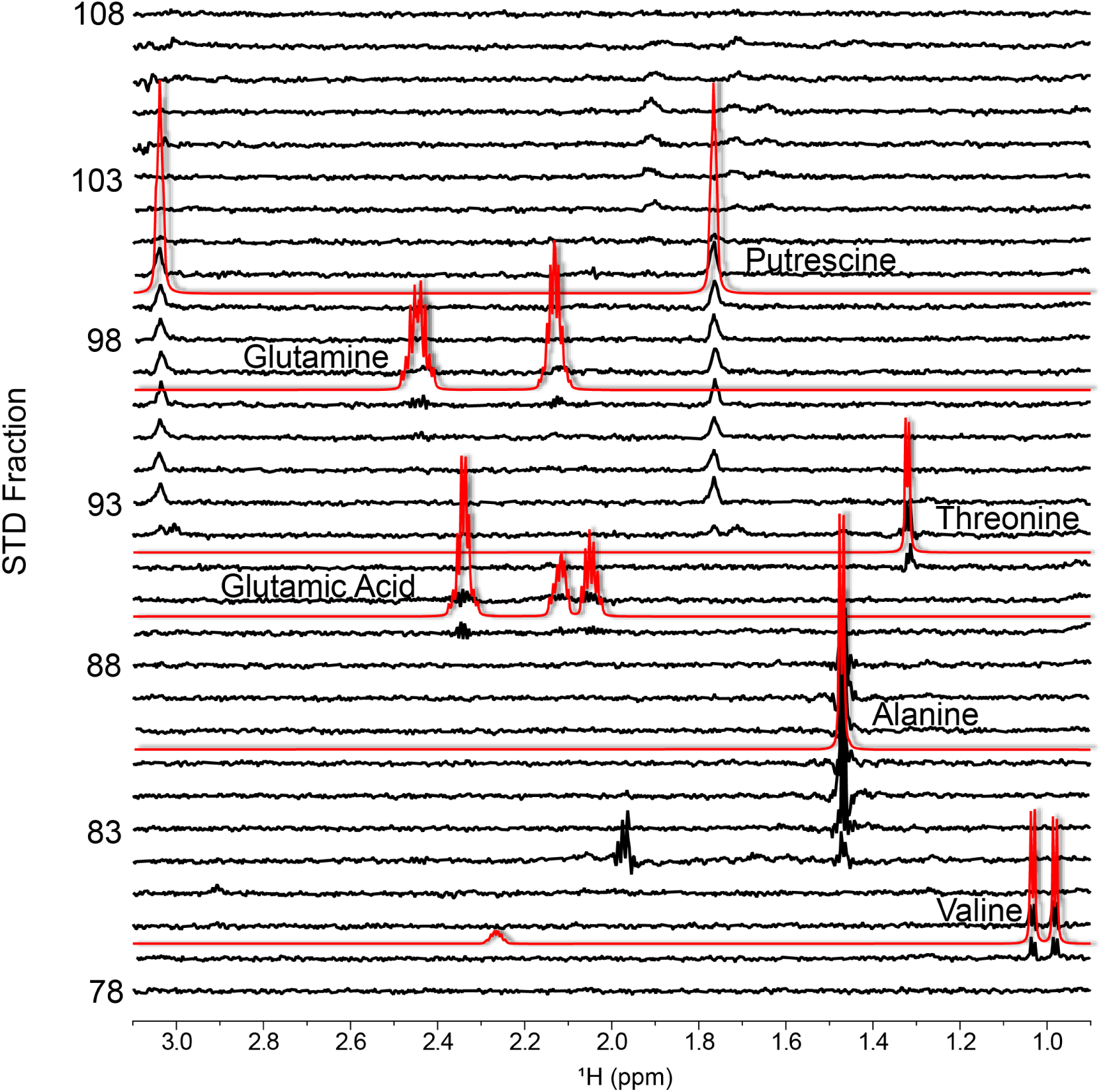
| STD NMR screen of the *C. elegans* mFL. STD NMR difference spectra (black) for library fractions overlaid with 1D ^1^H spectra of basis elements (red).

**SI Figure 4.**
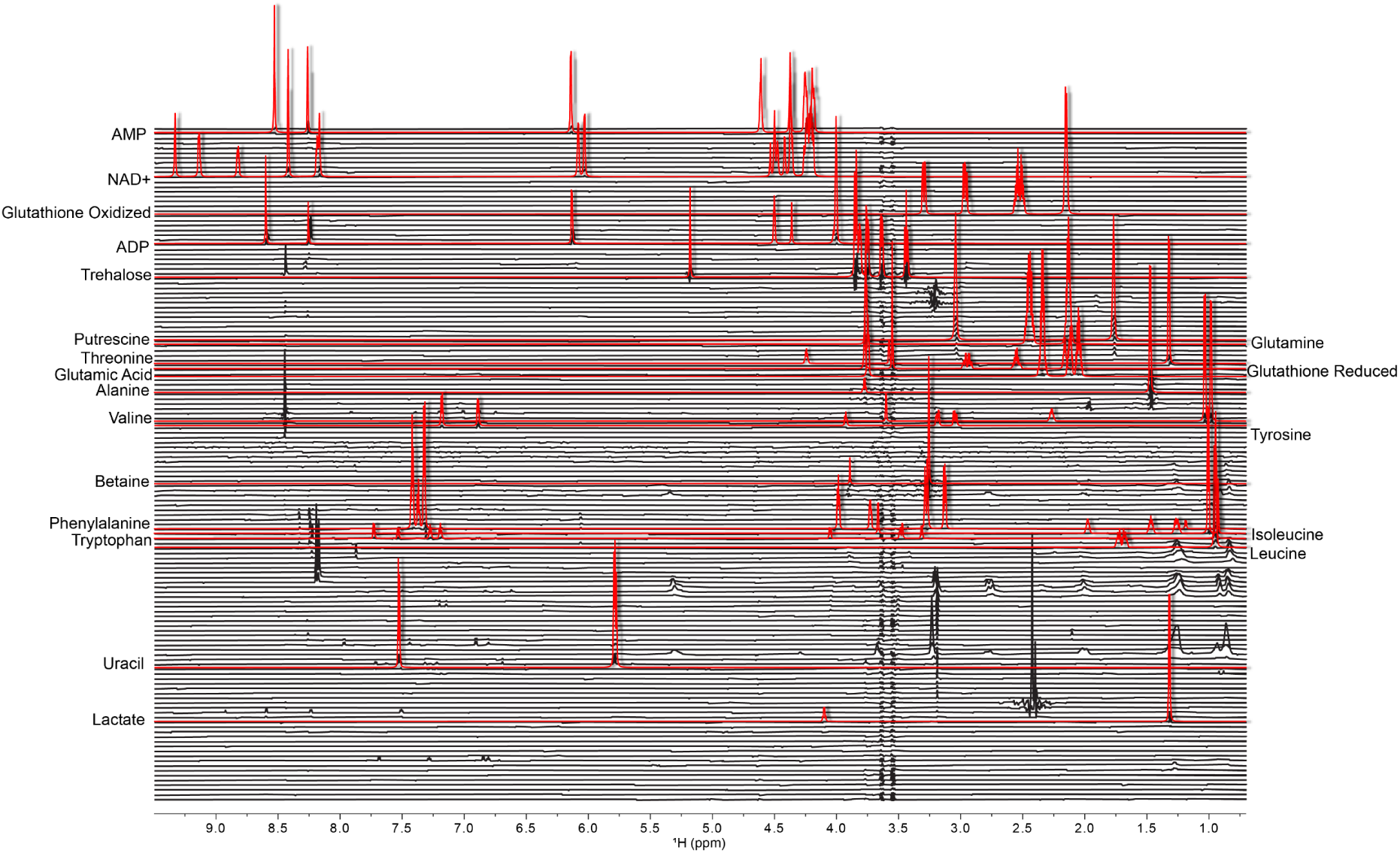
| STD NMR screen of the *C. elegans* mFL. STD NMR difference spectra (black) for every fraction in the library overlaid with 1D ^1^H spectra of basis elements (red).

**SI Figure 5.**
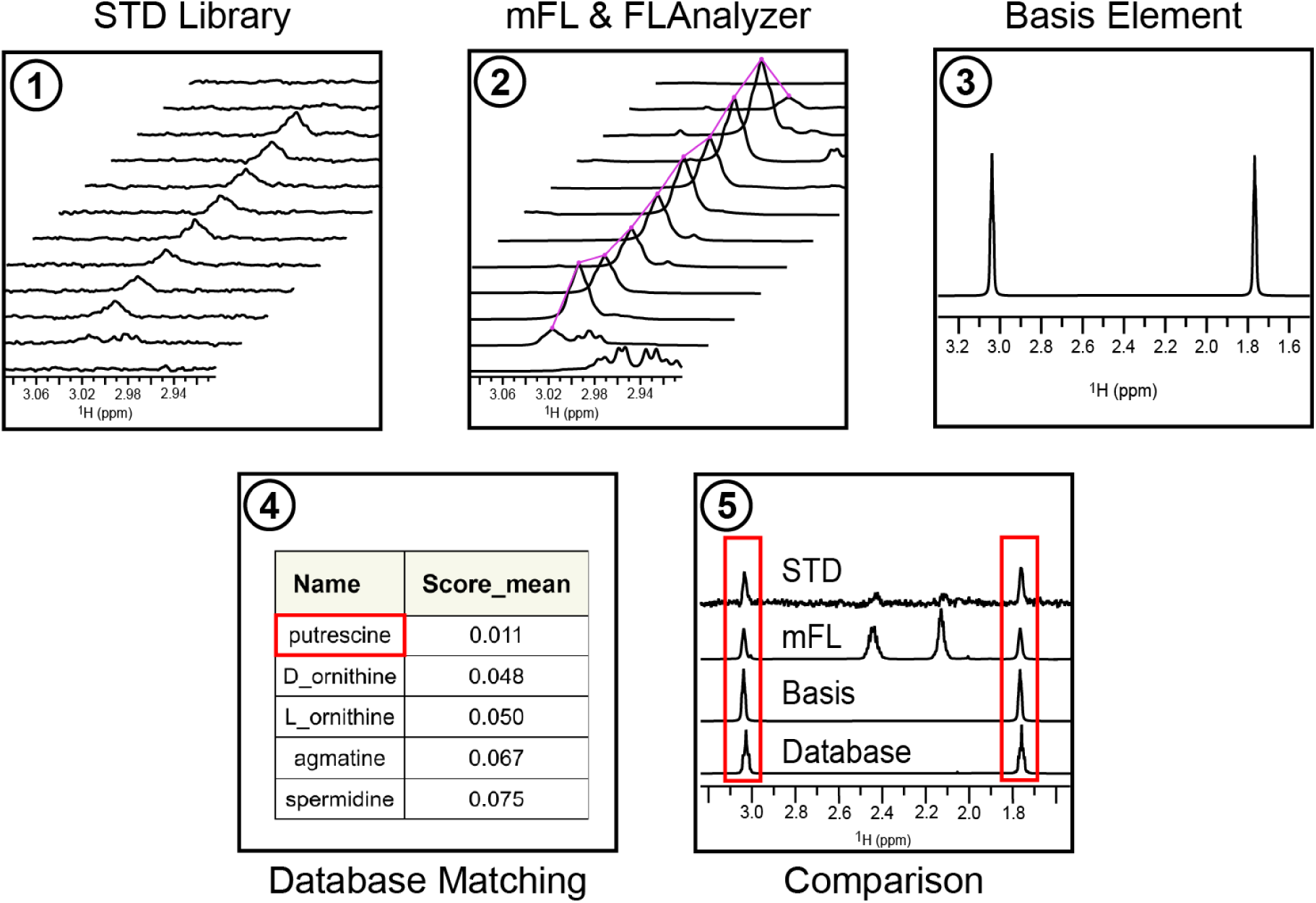
| Targeted extraction and identification of basis set elements. 1. An STD peak from a binder is visually identified in the STD library. 2. The same peak is identified in FLAnalyzer and traced as a driver peak. 3. A basis element that includes driver correlates is generated for the peak. 4. Database matching of the basis element identifies a candidate. 5. A synthetic database spectrum is compared to the basis element along with the original mFL and STD library data.

**SI Table 1.** – Non-targeted database match validation confidence (Uk = Unknown, NA = Not applicable, ND = Not determined)

| Basis # | Compound | BMRB Accession | Confidence |
|---|---|---|---|
| nt_basis_1 | Uk 1 | NA | ND |
| nt_basis_2 | Uk 2 | NA | ND |
| nt_basis_3 | Uk 3 | NA | ND |
| nt_basis_4 | betaine | bmse000069 | High |
| nt_basis_5 | trehalose | bmse000125 | High |
| nt_basis_6 | alanine | bmse000236 | High |
| nt_basis_7 | Uk 4 | NA | ND |
| nt_basis_8 | carnosine | bmse000246 | ND |
| nt_basis_9 | carnitine | bmse000211 | High |
| nt_basis_10 | glutathione_oxidized | bmse000170 | Medium |
| nt_basis_11 | Uk 5 | NA | ND |
| nt_basis_12 | choline | bmse000285 | High |
| nt_basis_13 | glutamic_acid | bmse000037 | High |
| nt_basis_14 | gamma_galactonolactone | bmse000141 | Medium |
| nt_basis_15 | threonine | bmse000049 | High |
| nt_basis_16 | arginine | bmse000711 | High |
| nt_basis_17 | Uk 6 | NA | ND |
| nt_basis_18 | diethanolamine | bmse000371 | Medium |
| nt_basis_19 | lactate | bmse000269 | High |
| nt_basis_20 | inosine_5_monophosphate | bmse000175 | Medium |
| nt_basis_21 | leucine | bmse000042 | High |
| nt_basis_22 | trimethylamine_N_oxide | bmse000426 | High |
| nt_basis_23 | glutamine | bmse000038 | High |
| nt_basis_24 | putrescine | bmse000109 | High |
| nt_basis_25 | valine | bmse000052 | High |
| nt_basis_26 | maltose | bmse000017 | Medium |
| nt_basis_27 | Uk 7 | NA | ND |
| nt_basis_28 | cyclohexanone | bmse000405 | Medium |
| nt_basis_29 | Uk 8 | NA | ND |
| nt_basis_30 | isoleucine | bmse000041 | High |
| nt_basis_31 | Uk 9 | NA | ND |
| nt_basis_32 | Uk 10 | NA | ND |
| nt_basis_33 | Uk 11 | NA | ND |
| nt_basis_34 | Uk 12 | NA | ND |
| nt_basis_35 | Uk 13 | NA | ND |
| nt_basis_36 | Uk 14 | NA | ND |
| nt_basis_37 | glycerol | bmse000856 | High |
| nt_basis_38 | Uk 15 | NA | ND |
| nt_basis_39 | pantothenate | bmse000287 | High |
| nt_basis_40 | Uk 16 | NA | ND |
| nt_basis_41 | Uk 17 | NA | ND |
| nt_basis_42 | Uk 18 | NA | ND |
| nt_basis_43 | betaine | bmse000069 | High |
| nt_basis_44 | Uk 19 | NA | ND |
| nt_basis_45 | Uk 20 | NA | ND |
| nt_basis_46 | 3_iodo_tyrosine | bmse000337 | Medium |
| nt_basis_47 | Uk 21 | NA | ND |
| nt_basis_48 | diethanolamine | bmse000371 | Medium |
| nt_basis_49 | Uk 22 | NA | ND |
| nt_basis_50 | serine | bmse000222 | High |
| nt_basis_51 | Uk 23 | NA | ND |
| nt_basis_52 | homocitrulline | bmse000442 | Medium |
| nt_basis_53 | Uk 24 | NA | ND |
| nt_basis_54 | glutathione_oxidized | bmse000170 | High |
| nt_basis_55 | ADP | bmse000004 | Medium |
| nt_basis_56 | methionine | bmse000044 | High |
| nt_basis_57 | Uk 25 | NA | ND |
| nt_basis_58 | Uk 26 | NA | ND |
| nt_basis_59 | proline | bmse000047 | High |
| nt_basis_60 | Uk 27 | NA | ND |
| nt_basis_61 | Uk 28 | NA | ND |
| nt_basis_62 | gamma_aminobutyric_acid | bmse000340 | High |
| nt_basis_63 | phenylalanine | bmse000045 | Medium |
| nt_basis_64 | Uk 29 | NA | ND |
| nt_basis_65 | Uk 30 | NA | ND |
| nt_basis_66 | Uk 31 | NA | ND |
| nt_basis_67 | Uk 32 | NA | ND |
| nt_basis_68 | Uk 33 | NA | ND |
| nt_basis_69 | Uk 34 | NA | ND |
| nt_basis_70 | Uk 35 | NA | ND |
| nt_basis_71 | Uk 36 | NA | ND |
| nt_basis_72 | Uk 37 | NA | ND |
| nt_basis_73 | Uk 38 | NA | ND |
| nt_basis_74 | Uk 39 | NA | ND |
| nt_basis_75 | Uk 40 | NA | ND |
| nt_basis_76 | Uk 41 | NA | ND |
| nt_basis_77 | alpha_glycerol_phosphate | bmse000193 | High |
| nt_basis_78 | Uk 42 | NA | ND |
| nt_basis_79 | Uk 43 | NA | ND |
| nt_basis_80 | Uk 44 | NA | ND |
| nt_basis_81 | Uk 45 | NA | ND |
| nt_basis_82 | linoleic_acid | bmse000604 | Medium |
| nt_basis_83 | NAD | bmse000053 | High |
| nt_basis_84 | Uk 46 | NA | ND |
| nt_basis_85 | carnitine | bmse000211 | High |
| nt_basis_86 | fructose_6_phosphate | bmse000012 | ND |
| nt_basis_87 | ethanolamine | bmse000276 | High |
| nt_basis_88 | Uk 47 | NA | ND |
| nt_basis_89 | spermidine | bmse000116 | High |
| nt_basis_90 | Uk 48 | NA | ND |
| nt_basis_91 | kynurenine | bmse000172 | Medium |
| nt_basis_92 | alpha_epsilon_diaminopimelic_acid | bmse000401 | High |
| nt_basis_93 | Uk 49 | NA | ND |
| nt_basis_94 | Uk 50 | NA | ND |
| nt_basis_95 | Uk 51 | NA | ND |
| nt_basis_96 | hypoxanthine | bmse000094 | Medium |
| nt_basis_97 | Uk 52 | NA | ND |
| nt_basis_98 | Uk 53 | NA | ND |
| nt_basis_99 | Uk 54 | NA | ND |
| nt_basis_100 | methanol | bmse000294 | High |
| nt_basis_101 | Uk 55 | NA | ND |
| nt_basis_102 | Uk 56 | NA | ND |
| nt_basis_103 | Uk 57 | NA | ND |
| nt_basis_104 | Uk 58 | NA | ND |
| nt_basis_105 | Uk 59 | NA | ND |

**SI Table 2.**
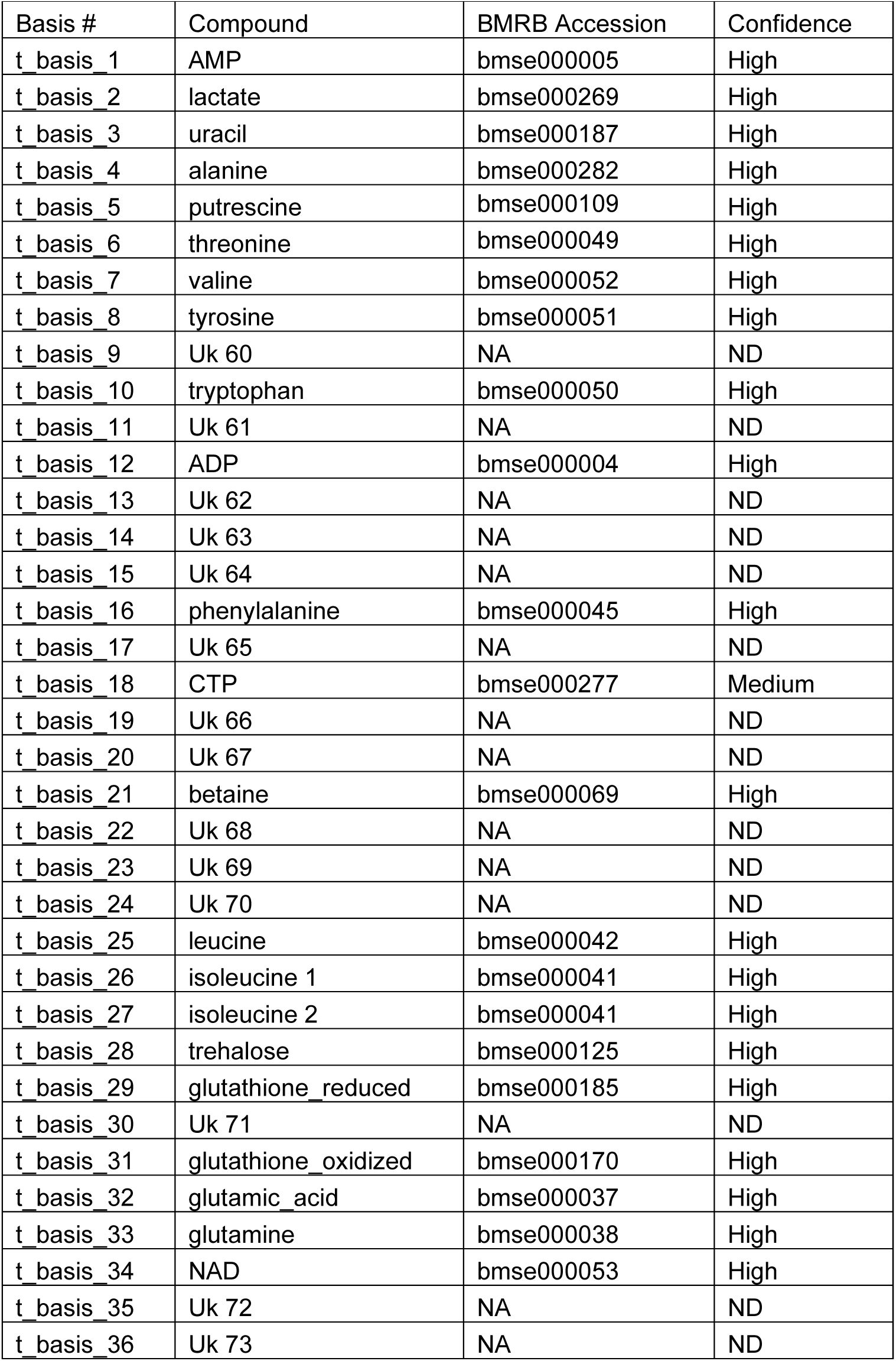
– Targeted database match validation confidence (Uk = Unknown, NA = Not applicable, ND = Not determined)

**SI Table 3.** – Cosine similarity of targeted and non-targeted mBS elements.

| Targeted Element | Best Non-targeted Element | Match Score |
|---|---|---|
| t_basis_5 | nt_basis_24 | 1 |
| t_basis_30 | nt_basis_22 | 1 |
| t_basis_21 | nt_basis_4 | 1 |
| t_basis_2 | nt_basis_19 | 1 |
| t_basis_7 | nt_basis_25 | 1 |
| t_basis_29 | nt_basis_10 | 0.999 |
| t_basis_27 | nt_basis_30 | 0.999 |
| t_basis_32 | nt_basis_13 | 0.996 |
| t_basis_33 | nt_basis_23 | 0.995 |
| t_basis_8 | nt_basis_46 | 0.995 |
| t_basis_18 | nt_basis_14 | 0.995 |
| t_basis_4 | nt_basis_6 | 0.987 |
| t_basis_25 | nt_basis_21 | 0.986 |
| t_basis_26 | nt_basis_45 | 0.981 |
| t_basis_6 | nt_basis_15 | 0.980 |
| t_basis_16 | nt_basis_41 | 0.975 |
| t_basis_19 | nt_basis_44 | 0.963 |
| t_basis_28 | nt_basis_5 | 0.957 |
| t_basis_24 | nt_basis_39 | 0.941 |
| t_basis_10 | nt_basis_97 | 0.923 |
| t_basis_34 | nt_basis_83 | 0.912 |
| t_basis_1 | nt_basis_20 | 0.858 |
| t_basis_14 | nt_basis_91 | 0.752 |
| t_basis_22 | nt_basis_48 | 0.752 |
| t_basis_3 | nt_basis_47 | 0.625 |
| t_basis_20 | nt_basis_23 | 0.430 |
| t_basis_35 | nt_basis_5 | 0.377 |
| t_basis_31 | nt_basis_54 | 0.375 |
| t_basis_12 | nt_basis_83 | 0.336 |
| t_basis_13 | nt_basis_71 | 0.297 |
| t_basis_23 | nt_basis_13 | 0.262 |
| t_basis_36 | nt_basis_51 | 0.213 |
| t_basis_9 | nt_basis_20 | 0.197 |
| t_basis_15 | nt_basis_42 | 0.156 |
| t_basis_11 | nt_basis_94 | 0.151 |
| t_basis_17 | nt_basis_101 | 0.028 |

